# Global climate change effect on Asian *Mus musculus*; Implication from last glacial maximum to the end of the 21^st^ century

**DOI:** 10.1101/2024.03.14.584992

**Authors:** Yaser Amir Afzali

## Abstract

Global climate change poses unprecedented challenges to biodiversity, prompting urgent investigations into its effects on various species. This study focuses on *Mus musculus*, a small rodent species and a crucial indicator of ecosystem health. Spanning from the last glacial maximum to the end of the 21^st^ century, employed Species Distribution Models (SDMs) to assess the impacts of climate change on *Mus musculus* and its four subspecies across Asia (*M. m musculus*, *M. m domesticus*, *M. m castaneus*, and *M. m bactrianus*). The SDMs reveal nuanced responses among subspecies, with *M. m. domesticus*, *M. m. musculus*, and *M. m. castaneus* facing potential habitat contractions, while *M. m. bactrianus* shows habitat expansion. Variable importance analysis highlights the significance of temperature-related variables, indicating the growing impact of rising temperatures on distribution patterns. Findings underscore the ecological implications of these shifts, emphasizing the need for tailored conservation strategies. The robustness of models, as indicated by high Area Under the Curve (AUC) values, enhances confidence in the reliability of predictions. Despite data limitations, this study contributes valuable insights into the complex dynamics between climate change and *Mus musculus* populations, guiding future conservation efforts in the face of ongoing global environmental transformations.

## INTRODUCTION

Over the past 150 years, the global average temperature of the Earth has risen by approximately 1.09 °C, a phenomenon attributed to anthropogenic activities (Hughes 2000; Wan et al. 2022). This rapid climate change poses a significant threat to ecosystems worldwide, with 16–33% of vertebrates currently facing global endangerment or extinction (Grimm et al. 2013; Dirzo et al. 2014). The resulting disruptions in natural ecosystems extend to modifications in function and structure, impacting species distribution, behavior, phenology, and inter-species interactions (Grimm et al. 2013; Lenoir & Svenning 2015; Rubenstein et al. 2019; Weiskopf et al. 2020; Amir Afzali et al. 2024). Climate change-induced alterations in terrestrial animal ranges are well-documented, with many species shifting towards higher latitudes or elevations (Darvish et al. 2012; Amir Afzali et al. 2017; Chen et al. 2011; Román-Palacios & Wiens 2020). Among these species, small rodents, constituting approximately 42% of mammalian diversity, serve as crucial indicators of ecosystem function due to their short lifespan, high reproductive capacity, and wide distribution (Amir Afzali & López-Antoñanzas 2024; Wan et al. 2022). Notably, studies by Levinsky et al. (2007) suggest that future climatic changes may lead to a substantial reduction in mammalian species richness in the Mediterranean region, with contrasting increases in the northeast and at higher elevations. To assess the impacts of climate change on biodiversity, particularly in the context of small rodent populations, Species Distribution Models (SDMs) emerge as invaluable tools (Araújo et al. 2005; Zhang et al. 2019; Yousefi et al. 2019). These models utilize species presence points and climatic data to predict species distributions and have been successfully employed in various contexts, including identifying future species distribution, climatic refugia, changes in distribution, and evaluating the effectiveness of protected areas under current and future climates (Ashrafzadeh et al. 2019; Hoveka et al. 2020; Prieto-Torres et al. 2020; Ramírez-Albores et al. 2020; Petersen et al. 2021; Sierra-Morales et al. 2021; Vaissi 2021). Previous studies have applied SDMs to assess the impacts of climate change on various rodent species (Cameron & Scheel 2001; Meserve et al. 2011; Jiang et al. 2013; Bean et al. 2014; Latinne et al. 2015; Gutiérrez-Tapia and Palma 2016; Austrich et al. 2021; Shiels et al. 2022; Wan et al. 2022). However, there is a notable gap in our understanding concerning the impacts of climate change on *Mus musculus*, specifically. Therefore, this study employs SDMs to predict the effects of climate change on the distribution of Mus musculus and its four subspecies across Asia (*M. m musculus* Linnaeus, 1758; *M. m domesticus* Schwarz and Schwarz, 1943; *M. m castaneus* Waterhouse, 1843; and *M. m bactrianus* Blyth, 1846). As this study delve into this investigation, it is essential to consider the potential cascading effects of climate change on *Mus musculus* and its subspecies. Small rodents play a crucial role in ecological processes, including seed dispersal, predation, and as prey for larger predators (Amir Afzali et al. 2018; Hulme-Beaman et al. 2019). The intricate web of ecological interactions involving *Mus musculus* makes it imperative to understand how changes in their distribution might reverberate throughout the ecosystem. Moreover, exploring the historical context of *Mus musculus*’s response to past climate fluctuations, such as during the last glacial maximum, can provide valuable insights into their adaptive capacity. Investigating the species’ resilience to climatic shifts over evolutionary timescales aids in predicting their potential responses to the rapidly changing climate of the 21^st^ century. This study not only addresses the current gaps in our knowledge but also lays the groundwork for understanding the complex dynamics between climate change and *Mus musculus* populations. By examining the species’ distribution over a vast temporal range, from the last glacial maximum to the present and projecting into the future, this study aimed to contribute to a comprehensive understanding of the challenges and opportunities these small mammals face in the wake of global climate change.

## MATERIALS AND METHODS

### Study area and occurrence points

Occurrence data for *Mus musculus* and its subspecies were sourced from the Global Biodiversity Information Facility (GBIF.org, 2023), providing a comprehensive dataset for the study. Additionally, incorporated opportunistic observations made by the author, enriching the dataset with localized insights. Upon compilation, the collected data underwent a rigorous processing phase to ensure accuracy and reliability. The distributions of *M. musculus* and each subspecies were meticulously mapped individually, employing Geographic Information System (GIS) techniques to visualize and analyze spatial patterns (Fig. 1). To maintain data integrity, conducted a thorough examination, removing outliers and duplicates that might compromise the precision of our analysis. This process involved the careful scrutiny of spatial coordinates and cross-referencing against existing literature and taxonomic databases. Following the quality control measures, our dataset comprised 340 records for *Mus musculus*, 82 records for *M. m. domesticus* (DOM), 32 records for *M. m. musculus* (MUS), 205 records for *M. m. castaneus* (CAS), and 21 records for *M. m. bactrianus* (BAC). Each record represents a crucial data point contributing to the understanding of the distributional dynamics of *Mus musculus* and its subspecies across the studied region. The spatial distribution map (Fig. 1) provides a visual representation of the geographic spread of *M. musculus* and its subspecies within the study area, offering insights into their ecological preferences and potential habitat associations. This robust dataset forms the foundation for our Species Distribution Models (SDMs), enabling us to predict and analyze the potential impacts of climate change on the distribution of *Mus musculus* across Asia.

**Figure 1.**
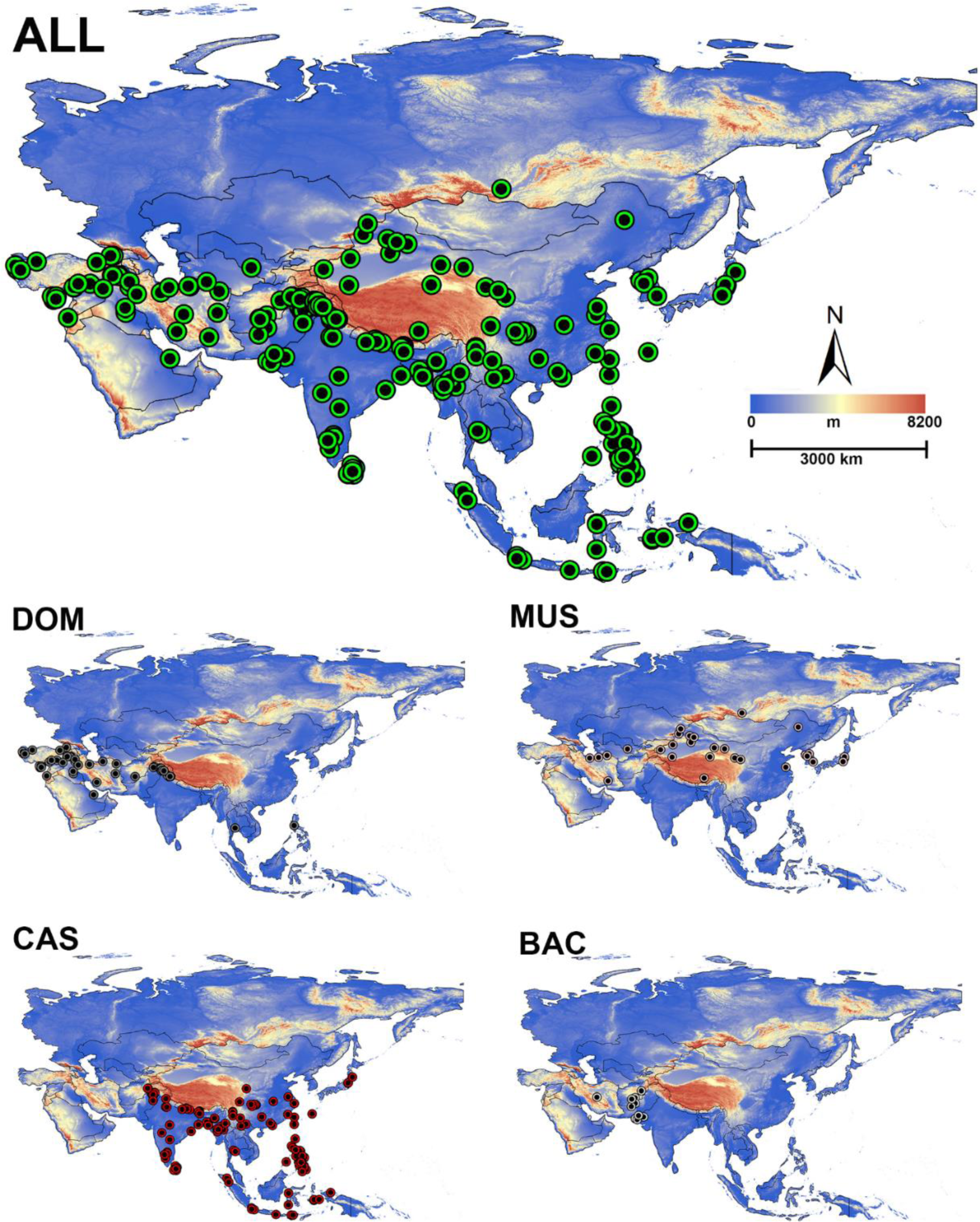
Spatial distribution map illustrating the geographic ranges of *Mus musculus* and its subspecies; *M. musculus* (ALL), *M. m. domesticus* (DOM), *M. m. musculus* (MUS), *M. m. castaneus* (CAS), and *M. m. bactrianus* (BAC) across the Asian continent.

### Climatic variables

Nineteen bioclimatic variables were obtained from the WorldClim series (Hijmans et al. 2005; http://www.worldclim.org) to characterize the environmental conditions for our species distribution models under Last Glacial Maximum (LGM), Current Condition (CC) and Future projection (F). CCSM4 used as general circulation model for the last glacial maximum and Future projection. For year 2070 (Future projection) considered high greenhouse gas emission scenarios, representative concentration pathway RCP 8.5 (Van Vuuren et al. 2011). As Figure 2 shows annual temperature predicted to increase in the future across the study area.

**Figure 2.**
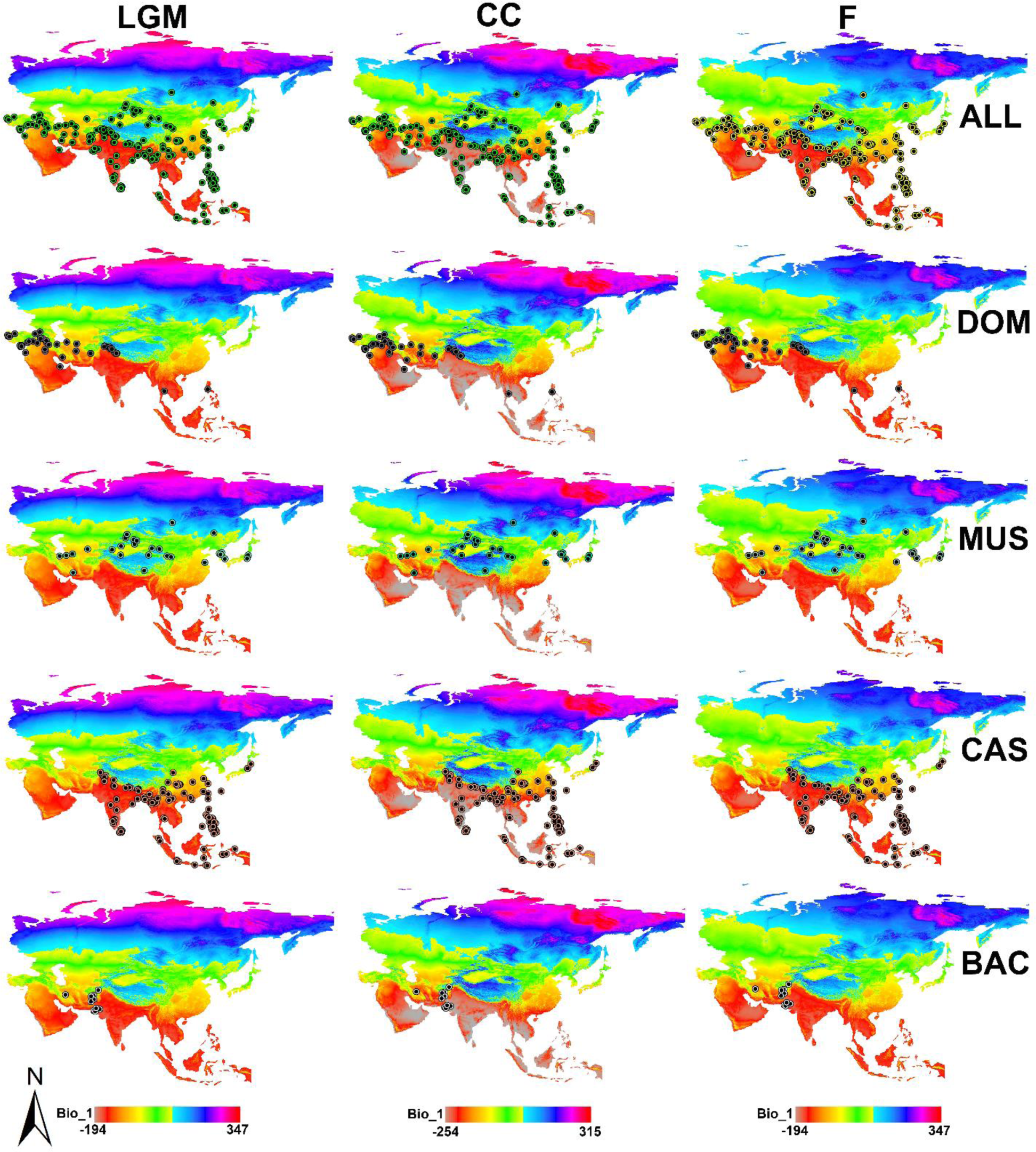
Predicted changes in annual temperature across the study area during three key periods; Last Glacial Maximum (LGM), Current Conditions (CC), and Future projection (F) along with occurrence records of *Mus musculus* (ALL) and its subspecies, including *M. m. domesticus* (DOM), *M. m. musculus* (MUS), *M. m. castaneus* (CAS), and *M. m. bactrianus* (BAC).

### Species distribution modeling

Maxent 3.4.4 (Phillips et al. 2006) utilized to estimate the probability of species occurrence. To perform this analysis, presence data used in conjunction with environmental variables, following the methodology outlined by Elith et al. (2006), Hernandez et al. (2006), and Rhoden et al. (2017). The models ran for three temporal conditions: the last glacial maximum (LGM∼22000 years BP), current conditions (CC∼1960-1990) and Future projection (F∼2061-2080). The Community Climate System Model (CCSM4) used as the general atmospheric circulation model. The spatial resolution was set at 2.5 arc-minutes (approximately 5 km^2^) for LGM and 30 arc-seconds (approximately 1 km^2^) for CC and F. In model training, %75 of the presence records was used, while %25 was randomly selected for testing. The procedure was repeated 15 times, with 5,000 iterations performed. The accuracy of the models was evaluated using Receiver Operating Characteristic (ROC) analysis. The Area Under the Curve (AUC) derived from the ROC plot, ranging between 0 and 1, served as the key metric. A model with an AUC value greater than 0.75 was considered robust and acceptable, while AUC values below 0.5 indicated a random prediction (Elith et al. 2006). To determine the importance of each climatic variable in explaining the species distribution model, conducted a jackknife procedure, following the approach outlined by Sillero & Carretero (2012). This procedure allowed to identify the key environmental variables driving the observed distribution patterns.

## RESULTS

### Climate change models

The models generated in this study exhibited strong performance, as indicated by the Receiver Operating Characteristic (ROC) curve values (AUC) ranging from 0.85 to 0.96 (Fig. 4). Analyses reveal distinct trends in habitat suitability for Mus musculus subspecies. Specifically, the suitability of habitats for *M. musculus* (ALL), *M. m. domesticus* (DOM), and *M. m. castaneus* (CAS) is projected to decrease. In contrast, *M. m. musculus* (MUS) and *M. m. bactrianus* (BAC) are anticipated to gain new suitable habitats in response to climate change (Fig. 3).

**Figure 3.**
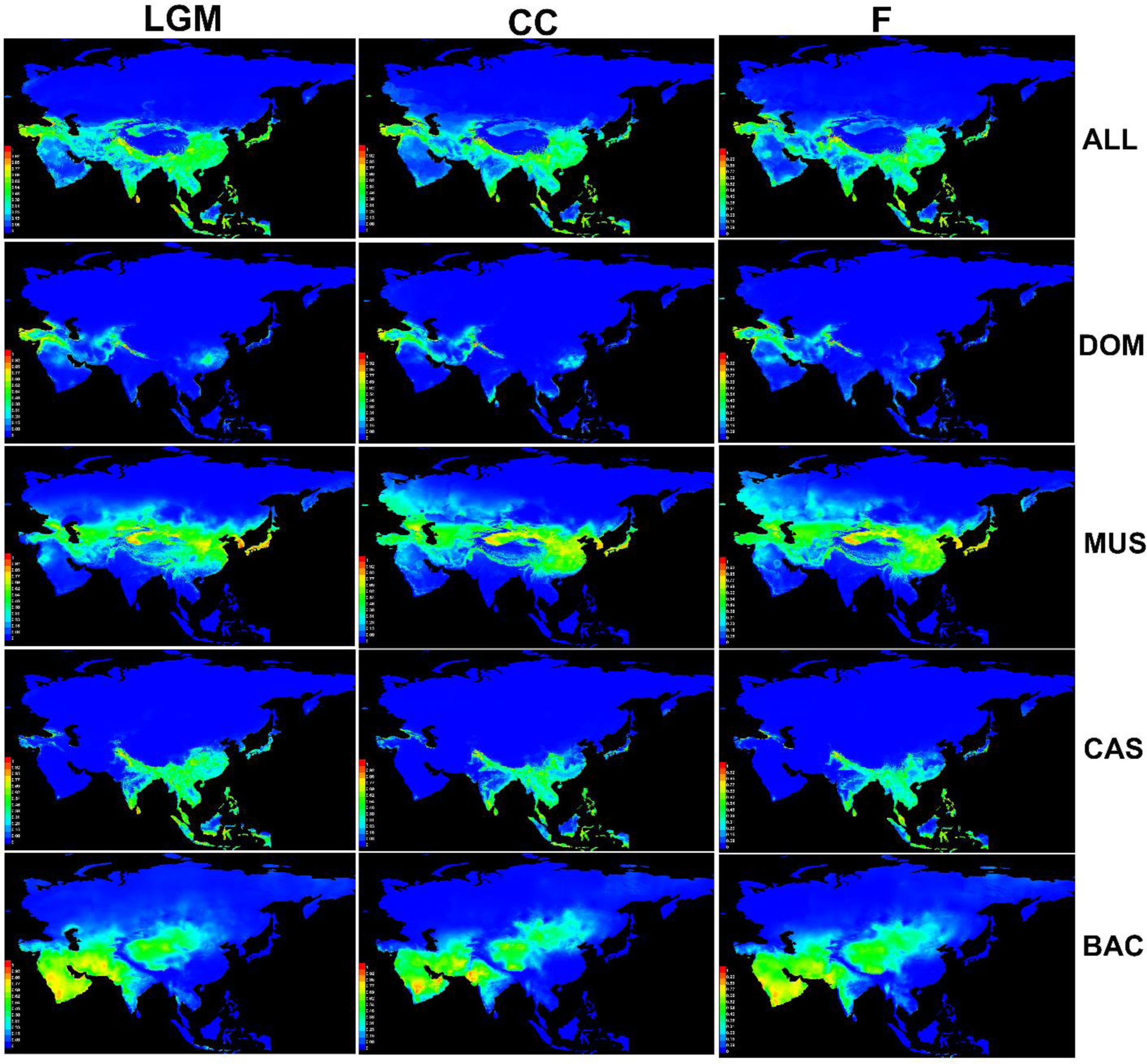
Species Distribution Models (SDMs) depicting the habitat suitability changes for *Mus musculus* (ALL) and its subspecies; *M. m. domesticus* (DOM), *M. m. musculus* (MUS), *M. m. castaneus* (CAS), and *M. m. bactrianus* (BAC) across three distinct temporal periods: Last Glacial Maximum (LGM), Current Conditions (CC), and Future projection (F).

**Figure 4.**
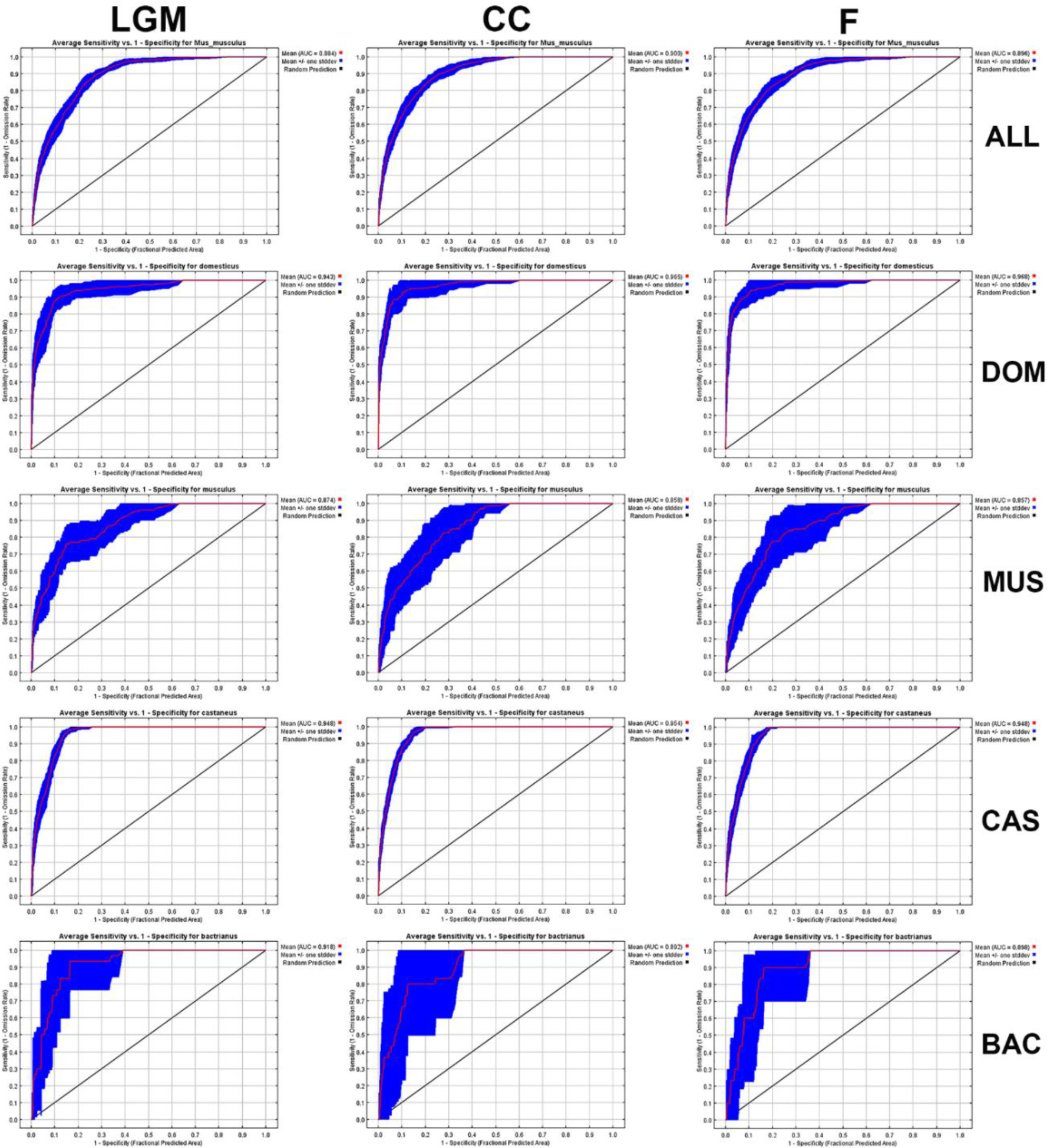
The Receiver Operating Characteristic (ROC) curve and AUC of *Mus musculus* and its subspecies; *M. musculus* (ALL), *M. m. domesticus* (DOM), *M. m. musculus* (MUS), *M. m. castaneus* (CAS), and *M. m. bactrianus* (BAC) during three distinct temporal periods: Last Glacial Maximum (LGM), Current Conditions (CC), and Future Projection (F).

### Species distribution modelling

In SDM analysis, models ran for the last glacial maximum (LGM), current conditions (CC), and future projection (F) through 15 replicates, all of which demonstrated high predictive power (Figs 3 & 4). The examination of habitat suitability for *Mus musculus* revealed dynamic changes across different temporal periods. *M. m. domesticus* (DOM) experienced a transition from LGM to F, exhibiting a reduction in suitable areas in East Asia and South India. However, there is a notable increase in habitat suitability in West Asia, suggesting potential shifts in ecological preferences under changing climatic conditions. *M. m. musculus* (MUS) displayed a significant gain in new suitable areas from LGM to F. The impact of global climate change is evident in the expansion of suitable habitats towards the North Pole and Northeast Asia. Although habitat suitability is anticipated to decrease in East Asia, particularly in Japan, the most suitable habitats persist in Central and East Asia. Conversely, *M. m. castaneus* (CAS) faced a decline in suitable areas, particularly in South and Southeast Asia, from LGM to F. These findings emphasize the vulnerability of certain areas that might no longer support the same level of habitat suitability for CAS in the face of changing climatic conditions. *M. m. bactrianus* (BAC) exhibited a unique pattern, showcasing more suitable habitats during the last glacial maximum compared to the current conditions. This suggests that BAC may have historically adapted to specific climatic conditions, and the contemporary climate might not fully align with its historical habitat preferences.

### Variable importance for house mouse

The assessment of variable importance in shaping the distribution of the house mouse (*Mus musculus*) is outlined in Table 1. Key environmental variables that significantly influenced the main models during different temporal periods are highlighted. During the last glacial maximum (LGM), the most influential variables included Bio_19 (Precipitation of Coldest Quarter), Bio_13 (Precipitation of Wettest Month), and Bio_2 (Mean Diurnal Range). These findings underscore the importance of precipitation-related variables during the historical period, indicating the crucial role of moisture conditions in shaping the habitat suitability of house mice. In current conditions (CC), the main influential variables shifted, with Bio_19 (Precipitation of Coldest Quarter), Bio_1 (Annual Mean Temperature), and Bio_15 (Precipitation Seasonality) taking precedence. The shift towards temperature-related variables emphasizes the contemporary significance of temperature patterns in determining the distribution of house mice in Asia. Looking into future projections (F), the most influential environmental variables were identified as Bio_11 (Mean Temperature of Coldest Quarter), Bio_3 (Isothermality), and Bio_1 (Annual Mean Temperature). The increasing importance of temperature-related variables in the future highlights the potential impact of ongoing climate change on shaping the distribution patterns of house mice, with temperature playing a central role. Comparatively, the transition from precipitation-related variables dominating during historical periods (LGM and CC) to temperature becoming more influential in future projections suggests a shift in the primary determinants of habitat suitability for house mice in Asia.

**Table 1.** Relative contributions of environmental variables to Maxent models, with emphasis on the top three influential variables for each model.

| Variables | Last Glacial Maximum |  |  |  |  | Current Condition |  |  |  |  | Future |  |  |  |  |
| --- | --- | --- | --- | --- | --- | --- | --- | --- | --- | --- | --- | --- | --- | --- | --- |
|  | Percent Contribution |  |  |  |  | Percent Contribution |  |  |  |  | Percent Contribution |  |  |  |  |
|  | DOM | MUS | CAS | BAC | ALL | DOM | MUS | CAS | BAC | ALL | DOM | MUS | CAS | BAC | ALL |
| Bio_1 (Annual Mean Temperature) | 0 | **20.9** | 0.5 | 0 | **17.3** | 0.7 | **40.7** | 1 | 0 | **36.7** | 0.3 | **30.1** | 0.4 | **0** | **28.7** |
| Bio_2 (Mean Diurnal Range) | 1.2 | 2 | 1.6 | **40.8** | 1.5 | 0.4 | 1.5 | 0.5 | **18.9** | 1.2 | 1 | 0.7 | 0.9 | **22** | 1 |
| Bio_3 (Isothermality (BIO2/BIO7) (×100)) | 0.4 | 0 | 4.3 | 0 | 2.8 | **22** | 1.8 | 4.1 | 0 | 5.2 | **33.7** | 5.3 | 3 | 4.2 | 5.8 |
| Bio_4 (Temperature Seasonality) | 5 | 4.9 | 1.5 | 6 | 6.1 | 6.8 | 3 | **25.9** | 0.1 | 6.4 | 1.8 | 0.5 | **21.3** | 0.4 | **9.4** |
| Bio_5 (Max Temperature of Warmest Month) | 0 | 1.1 | 0.9 | 0 | 0.9 | 0 | 0.4 | 1.1 | 0 | 1.4 | 0 | 0.1 | 0.9 | 0 | 2.1 |
| Bio_6 (Min Temperature of Coldest Month) | **13.5** | **26.5** | **16.5** | 0 | **33.6** | 1.5 | 0.5 | **11.8** | 0 | 0.5 | 1.8 | 1.6 | **19** | 0 | 1.9 |
| Bio_7 (Temperature Annual Range (BIO5-BIO6)) | 0.7 | 4.3 | 1.5 | 0 | 2.5 | 0.5 | 1 | 0.2 | 0 | 0.3 | 0.7 | 0 | 0.5 | 0 | 0.7 |
| Bio_8 (Mean Temperature of Wettest Quarter) | **10.9** | 1.8 | 1 | 0 | 1.3 | **7.7** | 2 | 0.9 | 0.1 | 1.8 | **19.5** | 2.7 | 0.9 | 0 | 2.3 |
| Bio_9 (Mean Temperature of Driest Quarter) | 3.7 | 2.8 | 0.4 | **33.7** | 0.5 | 0.5 | **4.1** | 2.2 | **24.8** | 1 | **16.3** | 3.1 | 1.5 | **28.5** | 1.5 |
| Bio_10 (Mean Temperature of Warmest Quarter) | 0.1 | 0.4 | 1.5 | 0.2 | 2.8 | 0.4 | 0.7 | 0.6 | 0 | 1 | 1.3 | 0.2 | 1.1 | 0 | 2.2 |
| Bio_11 (Mean Temperature of Coldest Quarter) | 7.6 | **23.6** | 1.4 | 1.2 | 3.7 | 0.5 | **32.6** | 3.8 | 0 | **16.3** | 1.3 | **41.6** | 5.2 | 0 | **21.9** |
| Bio_12 (Annual Precipitation) | 0.4 | 0.1 | 0.4 | **8.3** | 1.1 | 0.4 | 0.3 | 2.2 | 11.4 | 1.5 | 0 | 0.2 | 1.4 | 18.8 | 0.6 |
| Bio_13 (Precipitation of Wettest Month) | 0.7 | 4.5 | **55.3** | 0.3 | **8.7** | 0.4 | 0.5 | **33.7** | 1.5 | 10.8 | 8.8 | 2.5 | **27.2** | 0.2 | 8.8 |
| Bio_14 (Precipitation of Driest Month) | 0.1 | 0.4 | 0.5 | 0 | 0.3 | 0.2 | 1.1 | 1.6 | 0 | 0.5 | 0 | 1.4 | 0.7 | 3.1 | 0.5 |
| Bio_15 (Precipitation Seasonality) | 0.5 | 0.1 | 1.3 | 2.8 | 1.7 | 1.1 | 3 | 0.5 | **40.2** | 1 | 0.5 | **5.5** | 1.4 | **21.8** | 1.4 |
| Bio_16 (Precipitation of Wettest Quarter) | 2 | 0.1 | 1.4 | 0.7 | 4 | 0 | 0 | 1.4 | 0.3 | 1.5 | 0.1 | 0 | 4.4 | 0.7 | 2.7 |
| Bio_17 (Precipitation of Driest Quarter) | 0.6 | 0.5 | 2.2 | 1.6 | 2.4 | 3.4 | 0.9 | 3.2 | 0.2 | 1.3 | 8.1 | 1.5 | 4.6 | 0.1 | **7.7** |
| Bio_18 (Precipitation of Warmest Quarter) | 0.7 | 4.6 | **5.4** | 4.3 | 0.9 | 2.9 | 2.5 | 3.8 | 1.9 | 0.7 | 4.3 | 2.5 | 5.5 | 0.1 | 0.2 |
| Bio_19 (Precipitation of Coldest Quarter) | **52** | 1.2 | 2.1 | 0.1 | 7.8 | **50.5** | 3.5 | 1.5 | 0.5 | **10.9** | 0.5 | 0.3 | 0.2 | 0.1 | 0.6 |

## DISCUSSION

The findings of this study shed light on the potential impacts of global climate change on *Mus musculus* and its subspecies in Asia, spanning from the last glacial maximum to the end of the 21^st^ century. The utilization of Species Distribution Models (SDMs) based on a comprehensive dataset allowed us to examine the dynamic response of these small rodents to changing climatic conditions.

The examination of *Mus musculus* distribution during the last glacial maximum (LGM) provides valuable insights into the species’ historical response to climate fluctuations. The observed increase in suitable habitats for *M. m. musculus* (MUS) and *M. m. bactrianus* (BAC) during LGM suggests a capacity for adaptation and range expansion in response to colder climatic conditions. This adaptive potential aligns with the general understanding that small rodents often exhibit behavioral and physiological adaptations to survive in diverse environments. The study’s current condition (CC) models suggest that *M. m. domesticus* (DOM), *M. m. musculus* (MUS), and *M. m. castaneus* (CAS) face potential contractions in suitable habitats. This contraction is particularly evident in East Asia and South India for DOM, East Asia for CAS, and Japan for MUS. In contrast, *M. m. bactrianus* (BAC) shows an expansion of suitable habitats. These projections highlight the heterogeneity in responses among *Mus musculus* subspecies, emphasizing the need for nuanced conservation strategies.

The variable importance analysis underscores the role of temperature-related variables in shaping the distribution of *Mus musculus*. Annual Mean Temperature (Bio_1), Mean Diurnal Range (Bio_2), and Isothermality (Bio_3) were identified as key influencers during different periods. Notably, the increasing importance of temperature-related variables in future projections aligns with broader climate change patterns, suggesting that rising temperatures might become a predominant factor influencing the distribution of *Mus musculus*. The anticipated changes in *Mus musculus* distribution have ecological implications for Asia. The potential decrease in suitable habitats for some subspecies may impact ecosystem dynamics, affecting seed dispersal, predation, and serving as prey for larger predators. Furthermore, the observed shifts in distribution patterns may lead to altered interspecies interactions, potentially influencing the broader ecological network in which *Mus musculus* participates.

The high AUC values obtained for all models indicate the robustness and reliability of SDMs. This suggests that the models are effective in predicting the potential distributional changes of *Mus musculus* under different climatic scenarios. Despite the rigor in data processing and quality control, limitations exist in the availability of occurrence data. Incorporating more fine-scale data and continuous monitoring efforts could enhance the precision of future SDMs. Additionally, the reliance on SDMs assumes that species’ responses are solely climate-driven, not accounting for potential anthropogenic influences or interactions with other species.

Given the varied responses of *Mus musculus* subspecies to climate change, conservation efforts should adopt a differential approach. Subspecies facing habitat contractions might require targeted conservation strategies, including habitat restoration, while those experiencing habitat expansions may need monitoring to manage potential overpopulation and associated ecological impacts. Identifying areas with relatively stable conditions, such as potential climatic refugia, becomes crucial for conservation planning. Conservation efforts focused on protecting these refugial areas could serve as a strategy to safeguard *Mus musculus* populations in the face of changing climates.

## Conclusion

This study contributes to the understanding of how *Mus musculus* and its subspecies may respond to climate change in Asia, offering insights into historical adaptations and future projections. The nuanced examination of different subspecies and their responses underscores the complexity of climate change impacts on small mammal populations. As we move forward, it is imperative to integrate these findings into conservation practices and policy decisions, recognizing the importance of preserving biodiversity in the face of ongoing global climate change.

## REFERENCES

Araujo, M. B., Pearson, R. G., Thuiller, W., Erhard, M. (2005) Validation of species–climate impact models under climate change. Global change biology, 11, 1504–1513. 10.1111/j.1365-2486.2005.01000.x.

Amir Afzali, Y., Darvish, J., Yazdani Moghaddam, F. (2017) Study of rodents’ fauna of the Jiroft, Kerman Province in southeast of Iran. Iranian Journal of Biosystematics, 13, 119–129. 10.22067/ijab.v13i1.59907.

Amir Afzali, Y., Yazdani Moghaddam, F., Dianat, M., Mahmudi, A. (2018) Biosystematics Study of *Golunda ellioti* Gray, 1837 (Rodentia: Muridae) From Jiroft and Anbarabad Townships in Southeast of Iran. Journal of Research in Biology, 1, 1–5. 10.21859/jresbiol-e1522.

Amir Afzali, Y. & López-Antoñanzas, R. (2024) Molecular phylogeny and historical biogeography of Iranian murids (Rodentia: Muridae). Mammalian Biology, 104, 79–89. 10.1007/s42991-023-00390-3.

Amir Afzali, Y., Naderloo, R., Keikhosravi, A., Klaus, S. (2024) Geographic differentiation in the freshwater crab *Potamon persicum* Pretzmann, 1962 (Decapoda: Potamidae) in the Zagros Mountains; evidence from morphometry. Zoosystema, 46, 77–93. 10.5252/zoosystema2024v46a4.

Ashrafzadeh, M. R., Naghipour, A. A., Haidarian, M., Kusza, S., Pilliod, D. S. (2019) Effects of climate change on habitat and connectivity for populations of a vulnerable, endemic salamander in Iran. Global Ecology and Conservation, 19, e00637. 10.1016/j.gecco.2019.e00637.

Austrich, A., Kittlein, M. J., Mora, M. S., Mapelli, F. J. (2021) Potential distribution models from two highly endemic species of subterranean rodents of Argentina: which environmental variables have better performance in highly specialized species? Mammalian Biology, 101, 503–519. 10.1007/s42991-021-00150-1.

Bean, W. T., Prugh, L. R., Stafford, R., Butterfield, H. S., Westphal, M., Brashares, J. S. (2014) Species distribution models of an endangered rodent offer conflicting measures of habitat quality at multiple scales. Journal of Applied Ecology, 51, 1116–1125. 10.1111/1365-2664.12281.

Cameron, G. N. & Scheel, D. (2001) Getting warmer: effect of global climate change on distribution of rodents in Texas. Journal of Mammalogy, 82, 652–680. 10.1644/1545-1542(2001)082%3C0652:GWEOGC%3E2.0.CO;2.

Chen, I. C., Hill, J. K., Ohlemüller, R., Roy, D. B., Thomas, C. D. (2011) Rapid range shifts of species associated with high levels of climate warming. Science, 333, 1024–1026. 10.1126/science.1206432.

Darvish, J., Amir Afzali, Y., Hamidi, K. (2012) Further record of *Golunda ellioti* Gray, 1837 from South East of Iran with notes on its postcranial skeleton. Iranian Journal of Biosystematics, 8, 79–82. 10.22067/ijab.v8i1.25574.

Dirzo, R., Young, H. S., Galetti, M., Ceballos, G., Isaac, N. J., Collen, B. (2014) Defaunation in the Anthropocene. Science, 345, 401–406. 10.1126/science.1251817.

Grimm, N. B., Chapin III, F. S., Bierwagen, B., Gonzalez, P., Groffman, P. M., Luo, Y., Melton, F., Nadelhoffer, K., Pairis, A., Raymond, P. A., Schimel, J., Williamson, C. E. (2013) The impacts of climate change on ecosystem structure and function. Frontiers in Ecology and the Environment, 11, 474–482. 10.1890/120282.

Gutiérrez-Tapia, P. & Palma, R. E. (2016) Integrating phylogeography and species distribution models: cryptic distributional responses to past climate change in an endemic rodent from the central Chile hotspot. Diversity and distributions, 22, 638–650. 10.1111/ddi.12433.

Hoveka, L. N., van der Bank, M., Davies, T. J. (2020) Evaluating the performance of a protected area network in South Africa and its implications for megadiverse countries. Biological Conservation, 248, 108577. 10.1016/j.biocon.2020.108577.

Hughes, L. (2000) Biological consequences of global warming: is the signal already apparent? Trends in ecology & evolution, 15, 56–61. 10.1016/S0169-5347(99)01764-4.

Hulme-Beaman, A., Claude, J., Chaval, Y., Evin, A., Morand, S., Vigne, J. D., Dobney, K., Cucchi, T. (2019) Dental shape variation and phylogenetic signal in the Rattini tribe species of Mainland Southeast Asia. Journal of Mammalian Evolution, 26, 435–446. 10.1007/s10914-017-9423-8.

Jiang, G., Liu, J., Xu, L., Yu, G., He, H., Zhang, Z. (2013) Climate warming increases biodiversity of small rodents by favoring rare or less abundant species in a grassland ecosystem. Integrative Zoology, 8, 162–174. 10.1111/1749-4877.12027.

Latinne, A., Meynard, C. N., Herbreteau, V., Waengsothorn, S., Morand, S., Michaux, J. R. (2015) Influence of past and future climate changes on the distribution of three Southeast Asian murine rodents. Journal of Biogeography, 42, 1714–1726. 10.1111/jbi.12528.

Lenoir, J. & Svenning, J. C. (2015) Climate-related range shifts–a global multidimensional synthesis and new research directions. Ecography, 38, 15–28. 10.1111/ecog.00967.

Levinsky, I., Skov, F., Svenning, J. C., Rahbek, C. (2007) Potential impacts of climate change on the distributions and diversity patterns of European mammals. Biodiversity and Conservation, 16, 3803–3816. 10.1007/s10531-007-9181-7

Meserve, P. L., Kelt, D. A., Previtali, M. A., Milstead, W. B., Gutiérrez, J. R. (2011) Global climate change and small mammal populations in north-central Chile. Journal of Mammalogy, 92, 1223–1235. 10.1644/10-MAMM-S-267.1

Petersen, W. J., Savini, T., Gray, T. N., Baker-Whatton, M., Bisi, F., Chutipong, W., Cremonesi, G., Gale, G. A., Mohamad, S. W., Rayan, D. M., Seuaturien, N., Shwe, N. M., Siripattaranukul, K., Sribuarod, K., Steinmetz, R., Sukumal, N., Ngoprasert, D. (2021) Identifying conservation priorities for an understudied species in decline: Golden cats (*Catopuma temminckii*) in mainland Tropical Asia. Global Ecology and Conservation, 30, e01762. 10.1016/j.gecco.2021.e01762

Prieto-Torres, D. A., Lira-Noriega, A., Navarro-Sigüenza, A. G. (2020) Climate change promotes species loss and uneven modification of richness patterns in the avifauna associated to Neotropical seasonally dry forests. Perspectives in Ecology and Conservation, 18, 19–30. 10.1016/j.pecon.2020.01.002

Román-Palacios, C. & Wiens, J. J. (2020) Recent responses to climate change reveal the drivers of species extinction and survival. Proceedings of the National Academy of Sciences, 117, 4211–4217. 10.1073/pnas.1913007117

Ramírez-Albores, J. E., Prieto-Torres, D. A., Gordillo-Martínez, A., Sánchez-Ramos, L. E., Navarro-Sigüenza, A. G. (2021) Insights for protection of high species richness areas for the conservation of Mesoamerican endemic birds. Diversity and Distributions, 27, 18–33. 10.1111/ddi.13153

Rubenstein, M. A., Christophersen, R., Ransom, J. I. (2019) Trophic implications of a phenological paradigm shift: Bald eagles and salmon in a changing climate. Journal of Applied Ecology, 56, 769–778. 10.1111/1365-2664.13286

Shiels, A. B., Ramírez de Arellano, G. E., Shiels, L. (2022) Invasive rodent responses to experimental and natural hurricanes with implications for global climate change. Ecosphere, 13, e4307. 10.1002/ecs2.4307

Sierra-Morales, P., Rojas-Soto, O., Ríos-Muñoz, C. A., Ochoa-Ochoa, L. M., Flores-Rodríguez, P., Almazán-Núñez, R. C. (2021) Climate change projections suggest severe decreases in the geographic ranges of bird species restricted to Mexican humid mountain forests. Global Ecology and Conservation, 30, e01794. 10.1016/j.gecco.2021.e01794

Vaissi, S. (2021) Potential changes in the distributions of Near Eastern fire salamander (Salamandra infraimmaculata) in response to historical, recent and future climate change in the Near and Middle East: Implication for conservation and management. Global Ecology and Conservation, 29, e01730. 10.1016/j.gecco.2021.e01730

Van Vuuren, D. P., Edmonds, J., Kainuma, M., Riahi, K., Thomson, A., Hibbard, K., Hurtt, G. C., Kram, T., Krey, V., Lamarque, J. F., Masui, T., Meinshausen, M., Nakicenovic, N., Smith, S. J., Rose, S. K. (2011) The representative concentration pathways: an overview. Climatic change, 109, 5–31. 10.1007/s10584-011-0148-z

Wan, X., Yan, C., Wang, Z., Zhang, Z. (2022) Sustained population decline of rodents is linked to accelerated climate warming and human disturbance. BMC Ecology and Evolution, 22, 102. 10.1186/s12862-022-02056-z

Weiskopf, S. R., Rubenstein, M. A., Crozier, L. G., Gaichas, S., Griffis, R., Halofsky, J. E., Hyde, K. J. W., Morelli, T. L., Morisette, J. T., Muñoz, R. C., Pershing, A. J., Peterson, D. L., Poudel, R., Staudinger, M. D., Sutton-Grier, A. E., Thompson, L., Vose, J., Weltzin, J. F., Whyte, K. P. (2020) Climate change effects on biodiversity, ecosystems, ecosystem services, and natural resource management in the United States. Science of the Total Environment, 733, 137782. 10.1016/j.scitotenv.2020.137782

Yousefi, M., Kafash, A., Valizadegan, N., Ilanloo, S. S., Rajabizadeh, M., Malekoutikhah, S., Yousefkhani, S. S. H., Ashrafi, S. (2019) Climate change is a major problem for biodiversity conservation: A systematic review of recent studies in Iran. Contemporary Problems of Ecology, 12, 394–403. 10.1134/S1995425519040127

Zhang, Z., Xu, S., Capinha, C., Weterings, R., Gao, T. (2019) Using species distribution model to predict the impact of climate change on the potential distribution of Japanese whiting Sillago japonica. Ecological Indicators, 104, 333–340. 10.1016/j.ecolind.2019.05.023

